# Brief, repeated isoflurane anaesthesia elicits sex-dependent behavioural, cerebrovascular, and microglial alterations in mice

**DOI:** 10.64898/2026.02.11.705306

**Authors:** Zofia Sus, Ishan Verma, Shafiul H. Khan, Timofei Romanchuk, Ozgun Mavuk, Adina T. Michael-Titus, Simon McArthur, Jordi L. Tremoleda

**Affiliations:** Centre for Neuroscience, Surgery & Trauma, Blizard Institute, Faculty of Medicine & Dentistry, Queen Mary, University of London, 4, Newark Street, London E1 2AT, United Kingdom; Centre for Oral Immunobiology & Regenerative Medicine, Institute of Dentistry, Faculty of Medicine & Dentistry, Queen Mary, University of London, Blizard Institute, 4, Newark Street, London E1 2AT, United Kingdom

**Keywords:** anaesthesia, isoflurane, burrowing, nesting, neuroinflammation

## Abstract

General anaesthetics are indispensable tools in biomedical research, yet growing evidence shows that their effects on brain function extend beyond the transient loss of consciousness for which they are used. Although numerous preclinical studies have examined the impact of a single anaesthetic exposure, far less is known about the cumulative consequences of repeated administration protocols, despite their widespread use in longitudinal imaging, behavioural testing, and disease model studies. Here, we show that brief, repeated isoflurane exposures induce alterations in behaviour, cerebrovascular patterns, and microglial phenotype in adult CD1 mice, with several responses exhibiting pronounced sex specificity. Repeated anaesthesia suppressed locomotor activity selectively in females and impaired ethologically important maintenance behaviours, including nesting and burrowing, in both sexes. These behavioural effects were accompanied by sex-dependent changes in cerebral blood flow dynamics, with females displaying elevated cerebrovascular flow, yet males showing higher reactivity to isoflurane exposure. At the molecular level, repeated isoflurane exposure altered endothelial marker expression without modifying capillary density and shifted microglia toward a hyporeactive state. Together, these data reveal that even short, routine anaesthetic events produce biologically meaningful and sex-dependent effects on neural, vascular, and immune systems. These results indicate that the anaesthetic regimen is an often overlooked but influential factor that deserves careful consideration in pre-clinical neuroscience.

## Introduction

General anaesthetics, pharmacological agents that can reversibly induce a loss of consciousness, are amongst the most widely used medical drugs and are key enablers of a vast range of modern surgical and diagnostic procedures. While generally thought to be safely reversible and without extensive side effects, it is important to note that they have an acute impact upon the respiratory and cardiovascular systems (Kim et al., 2017), and more persistent actions upon the brain, sometimes lasting well after the recovery of consciousness (Wu et al., 2019). Critically, these effects may accumulate following repeated exposure to anaesthesia, as identified in neurodevelopmental studies (Robinson et al., 2022).

A range of neurocognitive impacts have been reported following general anaesthetic exposure, with post-operative delirium, the short term disruption to attention and consciousness, being particularly tied to these agents (Siddiqi et al., 2006; Maldonado, 2008; Wu et al., 2025). Moreover, such brain vulnerability may persist, when repeated exposure to general anaesthesia for invasive examinations leads to prolonged cognitive deficits, which can aggravate neurodegeneration (Peng et al., 2023; Alharbi et al., 2024). Clinical studies both support and refute links between general anaesthetic use and perioperative neurocognitive disorders (Belrose and Noppens, 2019; Neuman et al., 2021), but given the wide range of confounders, not least that general anaesthetics are rarely used without pre-existing medical issues, it is extremely challenging to determine to what degree the two are linked.

For this reason, a growing body of preclinical studies have begun to investigate the cognitive effects of general anaesthetic exposure, reviewed in (Guo et al., 2023). Several key findings are apparent from this scoping review, with most studies reporting cognitive deficits, particularly in older animals, typically in aspects of learning and memory, but also in attention. Motor deficits were generally not described, and effects tended to disappear within two weeks of anaesthetic exposure. Rates of deficit appearance were similar in studies of rats or mice, suggesting cross-species effects. Potential sex-dependent differences were generally not investigated, as most studies reported the use of only male animals.

A substantial majority of these pre-clinical studies have focused on the effect of a single exposure to anaesthetic (Guo et al., 2023), but it is not at all uncommon for animals to undergo repeated rounds of general anaesthesia within a relatively short time frame, for example as part of an *in vivo* serial imaging protocol. Despite the value of examining such longitudinal responses *in vivo* to assess the progression of neurodegeneration and/or therapeutic responses, the implications of repeated anaesthetic exposure upon cerebrovascular homeostasis, cognition and neuroinflammation as relevant confounders, are generally poorly considered. What evidence is available suggests that this is not just a hypothetical concern. For example, repeated isoflurane exposure has been shown to cause mild distress in mice, affecting food intake, and exploratory behaviours (Hohlbaum et al., 2017) and in guinea-pigs, triggering an hypotensive cardiovascular response (Schmitz et al., 2017), and to induce acute and persistent deficits to motor function (Bajwa et al., 2019) and cognitive performance in mice (Long II et al., 2016; Bajwa et al., 2019). Similarly, repeated, but not single, exposure to the chemically related general anaesthetics sevoflurane and desflurane have been shown to impair cognition in both mice (Shen et al., 2013) and rats (Callaway et al., 2015), respectively. Interestingly, some authors reported that male and female mice perceived the distress differently, with females showing a greater susceptibility to the effects of repeated anaesthesia (Hohlbaum et al., 2017; Wasilczuk et al., 2024).

These cognitive and motor deficits are likely to confound any behavioural outcomes in central nervous disease (CNS) models, thus it is important to understand the consequences of repeated anaesthetic exposure, and the cell- and tissue-level changes that may underpin cumulative effects on brain homeostasis, neuroinflammation and model outcomes, not to mention the important implications they may have for animal care and welfare. We therefore sought to address this gap in understanding, by investigating the effects of repeated, brief induction of general anaesthesia with the inhalational agent isoflurane, widely used in preclinical modelling, upon key aspects of central nervous system physiology in wild-type mice. To this end, adult male and female CD1 mice were exposed to isoflurane on six, brief serial occasions, followed by home cage automatic behavioural assessment, and analysis of their cerebrovascular function and neuroinflammatory responses.

## Materials & Methods Ethical statement

All experimental procedures undertaken in this study were approved by the Queen Mary University of London (QMUL) Animal and Welfare and Ethical Review Body and were conducted under an approved UK Home Office Project Licence (PPL PP3042365), conforming to the UK Animals (Scientific Procedures) Act of 1986.

### Animal care and husbandry

Adult male and female CD-1 mice (12-14 weeks old; male average weight 38.15 ± 3.41 g, female average weight 30.98 ± 2.92 g at the start of the study; Charles River UK Ltd., Margate, UK; see individual studies for group size details) were housed in single sex groups of three in individual ventilated cages (floor area 1300 cm^2^; Allentown Europe, UK) with 1-1.5 cm wood chip bedding (Lignocel®, UK Ltd.), shredded nesting paper, sterilised hay, and environmental enrichment (three tunnels), at 21 ± 2°C, 55 ± 10 % relative humidity under a 12-hour light-dark cycle (lights on 06:00-18:00). Drinking water and standard chow (LabDiet EURodent 14% Diet, Land O’Lakes Inc., Missouri, U.S.A.) were provided *ad libitum*, and visual welfare checks were performed daily. Animals were allowed to acclimatise to the local environment for at least one week prior to data collection.

### Repeated isoflurane anaesthesia

Mice were exposed to a mix of 5% isoflurane, 95% oxygen (2 l/min flow) for 1-2 minutes to induce anaesthesia, followed upon loss of consciousness (defined as lack of a pedal reflex and slower breathing rate) with treatment for 5 minutes with 2.5 % isoflurane in oxygen (2 l/min flow rate), and recovery in normal air. Anaesthesia regime was selected to mimic approaches commonly used in serial imaging (e.g. optical imaging) experiments. Body temperature was maintained through use of a warming pad throughout. This procedure was repeated five further times, taking place on each Monday, Wednesday and Friday over two weeks. In each instance, procedures were started at 09:00 to minimise variations associated with circadian rhythm.

### Home cage behaviour analysis

At least one week prior to the repeated anaesthetic exposure procedure, an RFID transponder (BioTherm13 Passive Integrated Transponder; BioMark, Boise, ID, USA) was implanted subcutaneously in the lower abdominal flank of each mouse under brief isoflurane anaesthesia (4% isoflurane in oxygen, <3 minutes) as described previously (Yip et al., 2019). Automated behavioural recordings were taken using the ActualHCA™ home cage analyser system (Actual Analytics Ltd, UK) as reported previously (Yip et al., 2019). Briefly, continuous automated measures of RFID-identified mouse body temperature and horizontal position above the apparatus baseplate were collected alongside high-definition video recordings taken under infrared lighting (15-minute time bins). During data collection, human interaction was restricted to visual observation only, with no direct intervention to the cage. Baseline data (66 hours) was collected after acclimatization and one week before the first isoflurane dose. Post-exposure recordings (42 hours) were taken after each of the six isoflurane exposures.

Manual ethogram analysis of the video recordings was performed from 06:00-06:30 and 18:00-18:30 at baseline and within the first 24 h following isoflurane exposure, chosen as periods of consistent high locomotor activity. A modified version of the Stanford Mouse Behaviour ethogram (Graner) was used to assess exploratory, social and maintenance behaviours (Supplemental Table 1), with assessors blinded as to treatment group. Behaviours were analysed at the cage level, with n=3 mice per cage. Baseline recordings included n=3 cages per sex (n=6 total), whereas the repeated-anaesthesia conditions (ISO1–ISO6) included n=2 cages per sex (n=4 total). The incidence of behavioural patterns for n=3 animals per each cage was plotted. Each cage was assessed twice, once during the light phase (06:00–06:30) and once during the dark phase (18:00–18:30), yielding n=6 data points for baseline and n=4 data points for each ISO condition per time phase.

### *In vivo* brain perfusion imaging

Laser speckle perfusion imaging of the brain was performed using a moorFLPI Full-Field Laser Perfusion Imager (Moor Instruments, Devon, UK) according to manufacturer’s instructions and as described previously (Yip et al., 2019). Brain perfusion imaging data were acquired in two experimental cohorts: baseline (only subjected to isoflurane anaesthesia during the imaging procedure) and animals exposed to the six episodes of isoflurane anaesthesia, as described above (n=4 male and n=4 females per experimental cohort). Briefly, after induction of anaesthesia with isoflurane, a midline scalp incision was made to expose the skull. The CCD camera of the moorFLPI system was placed approximately 30 cm above the skull using an articulating arm, and video imaging at a display rate of 25 Hz and 4 ms exposure commenced. Three regions of interest (ROIs) were selected for each animal: two on either side of the superior sagittal sinus, avoiding large vessels, and one spanning the length of the sinus (Figure 3A). Each experimental session consisted of two repetitions of an isoflurane exposure protocol, where the isoflurane concentration was increased to 5% isoflurane in oxygen, followed by a return to 2% isoflurane in oxygen.

### Microglia extraction & analysis

Male and female mice (n=12 sham treated, n=6 exposed to ISO6; for each sex) were humanely killed by exsanguination by intracardiac perfusion with 0.1 M phosphate buffered saline (PBS; Merck Ltd., Gillingham, UK) under deep anaesthesia (isoflurane 5% in oxygen). Brains were rapidly removed and transferred to a collection buffer of PBS containing 15 mM HEPES, 0.5% D-glucose, 50 mg/mL penicillin-streptomycin (Thermofisher Scientific Ltd., Horsham, UK). Tissue was cut into ∼1 mm^3^ pieces and cells were dissociated by incubation for 30 minutes at 37°C in PBS containing 1.5 U/ml papain, 0.5 mg/ml deoxyribonuclease I, 5 mM L-cysteine, 5 mg/ml D-glucose, 0.4 % bovine serum albumin, 2 mM HEPES, 50 mg/ml penicillin-streptomycin (all Thermofisher Scientific, UK) accompanied by periodic gentle trituration. Following passage through a 40 μm cell strainer, myelin was removed by resuspension of the cells in 0.9 M sucrose in PBS (Merck, Germany) and centrifugation at 800 g for 10 minutes at 4°C. Cells were washed and resuspended in PBS at 4°C prior to immunostaining.

Cells were pre-incubated for 20 minutes with purified rat CD16/CD32 monoclonal antibody (0.5 mg/ml; Thermofisher Scientific Ltd., UK) to block surface Fcγ receptors, followed by incubation for 30 minutes at 4°C in darkness with a cocktail of PE conjugated anti-mouse CD45, APC Fire750-conjugated anti-mouse CD11b, PECy7-conjugated anti-mouse CD40, BV421-conjugated anti-mouse I-A/I-E (MHCII), BV605-conjugated anti-mouse CD68 and AF488-conjugated anti-mouse CD206, supplemented with HELIX NP NIR cell viability dye (all Biolegend Inc., San Diego, USA). Cells were analysed using an ACEA NovoCyte 3000 flow cytometer (Agilent Technologies Inc., Santa Clara, CA, USA) equipped with blue, red and violet lasers. A total of 20,000 events per sample were collected and data was analysed using NovoExpress 1.6.3 (Agilent Technologies Inc., USA), according to the gating strategy outlined in Supplemental Figure 1.

### Quantitative reverse-transcription polymerase chain reaction (qRT-PCR)

Animals (n=4 naive; n=12 exposed to ISO6) were humanely killed by exsanguination by intracardiac perfusion with 0.1 M PBS (Merck Ltd., Gillingham, UK) under deep anaesthesia (5% isoflurane in oxygen). Following perfusion, brains were removed from the skull and snap frozen in liquid nitrogen. Total RNA was prepared using TRIzol reagent (Life Technologies Ltd.) and then reverse-transcribed with Ultrascript 2.0 reverse transcriptase (PCRBiosystems Ltd., London, UK) according to the manufacturer’s protocols. Resultant cDNA was then analysed by real-time PCR in duplicate, using the Quantitect primer system (primer sets: *Cldn5* QT00254905, *Dlg4* QT00121695, *Il1b* QT01048355, *Il10* QT00106169, *Ocln* QT00111055, *Pecam1* QT01052044, *Syp* QT01042314, *Tjp1* QT00493899 and *Tnfa* QT00029162; all Qiagen Ltd.) and SyGreen (Blue) Lo-ROX master mix (PCRBiosystems Ltd., UK), according to conditions consisting of 95°C for 2 minutes, then 40 cycles of 95°C for 5 seconds, 55°C for 25 seconds, and 72°C for 20 seconds. Reactions were performed in 384-well format using a LightCycler 480 Instrument II (Roche Diagnostics Ltd., Burgess Hill, UK) and analysed with LightCycler 480 software v.1.5.1.62 (Roche Diagnostics Ltd., UK). Due to limited numbers of control animals, data were pooled by sex, and isoflurane-induced fold changes from control were calculated as 2^−ΔΔCt^.

### Cerebrovascular labelling

Male and female mice (n=4 controls, n=6 exposed to ISO6; for each sex) were humanely killed by exsanguination by intracardiac perfusion with 0.1 M phosphate buffered saline (PBS; Merck Ltd., Gillingham, UK) under deep anaesthesia (isoflurane 5% in oxygen). Following perfusion, brains were removed from the skull and immersed in 10% w/v formalin (Merck Ltd., UK) for 24 h post perfusion. Brains were then embedded in paraffin and cut coronally to 6 μm thickness with a rotary microtome (RM2255, Leica Microsystems, Milton Keynes, UK); sections were mounted on glass microscope slides. Following removal of paraffin and rehydration with xylene and ethanol, sections were subjected to antigen retrieval by incubation in 10 mM sodium citrate buffer (pH 6; Merck Ltd., UK) for 20 minutes at 90°C. Non-specific binding was blocked by incubation for 2 hours at room temperature with 10 % normal donkey serum, 0.025 % Triton x-100 in 0.1M Tris-buffered 0.9% saline (all Merck Ltd., UK) and sections were labelled with AF594-conjugated tomato lectin (Vector Laboratories, U.K. DL-1177) at 1:200 in 0.1 M Tris-buffered saline for 1 hour at 37°C. Hoechst 33342 (1:500, Merck Ltd., UK) was used to visualize nuclei. Image capture was performed at 20x using an IN-Cell Analyzer 2200 (GE Healthcare Life Sciences, USA). Quantitative analysis of the proportion of AF594-conjugated tomato lectin coverage in randomly selected cortical ROI was carried out using the HALO software (Indica Labs, Albuquerque, United States).

### Statistical analysis

Sample sizes were calculated to detect differences of 15% or more with a power of 0.85 and α set at 5%, calculations being informed by pilot data. Experimental data are expressed as mean ± SEM. Statistical analysis was performed using either GraphPad Prism (v10.2.0; GraphPad Software Inc., San Diego, CA) or RStudio 2022.07.01, with a P value of <0.05 being considered significant. Full blinding to treatment condition during sample processing, data collection and analysis was maintained. In general, following establishment of normality of distribution using the Shapiro – Wilk test, quantitative data was analysed with two-tailed Student’s *t*-tests for two groups or, for multiple comparisons, one-or two-way ANOVA with Tukey’s HSD *post hoc* test as appropriate. For HCA data, a mixed-model ANOVA was used to analyse repeated measures over time (15-min bins across the 42-hour periods) with sex as a fixed factor. This model accounted for missing values due to intermittent RFID signal loss (e.g., when animals were out of sensor range-e.g. hanging/ climbing). For semi-quantitative video analysis of behaviour, which was not normally distributed, a Mann-Whitney U-test was first used to compare baseline behaviours between sexes, followed by analysis of behavioural changes across all incidences of anaesthesia using a two-way ANOVA with Greenhouse-Geisser correction for unequal variance, followed by Dunnett’s multiple comparisons test (baseline vs each exposure stage).

## Results

### Sex-specific effects of anaesthesia upon locomotor activity and body temperature

We first examined the effects of repeated brief exposure to isoflurane anaesthesia upon murine CNS function by measuring diurnal and nocturnal locomotor behaviour and body temperature in RFID-identified CD-1 mice maintained within their home cages (Figure 1A-C). Statistical analysis revealed a significant effect of isoflurane exposure on locomotor distance in both light (F_1,20_=9.677, P=0.006) and dark phases (F_1,20_=17.749, P<0.001). We identified a notable sex difference in this parameter, with female mice showing a distinct reduction in mean distance travelled at the end of the sixth isoflurane exposure compared to their baseline activity in both light (P=0.004) and dark phases (P=0.005; Figure 1D), whereas male animals showed no significant anaesthesia-related effects on locomotor function.

**Figure 1:**
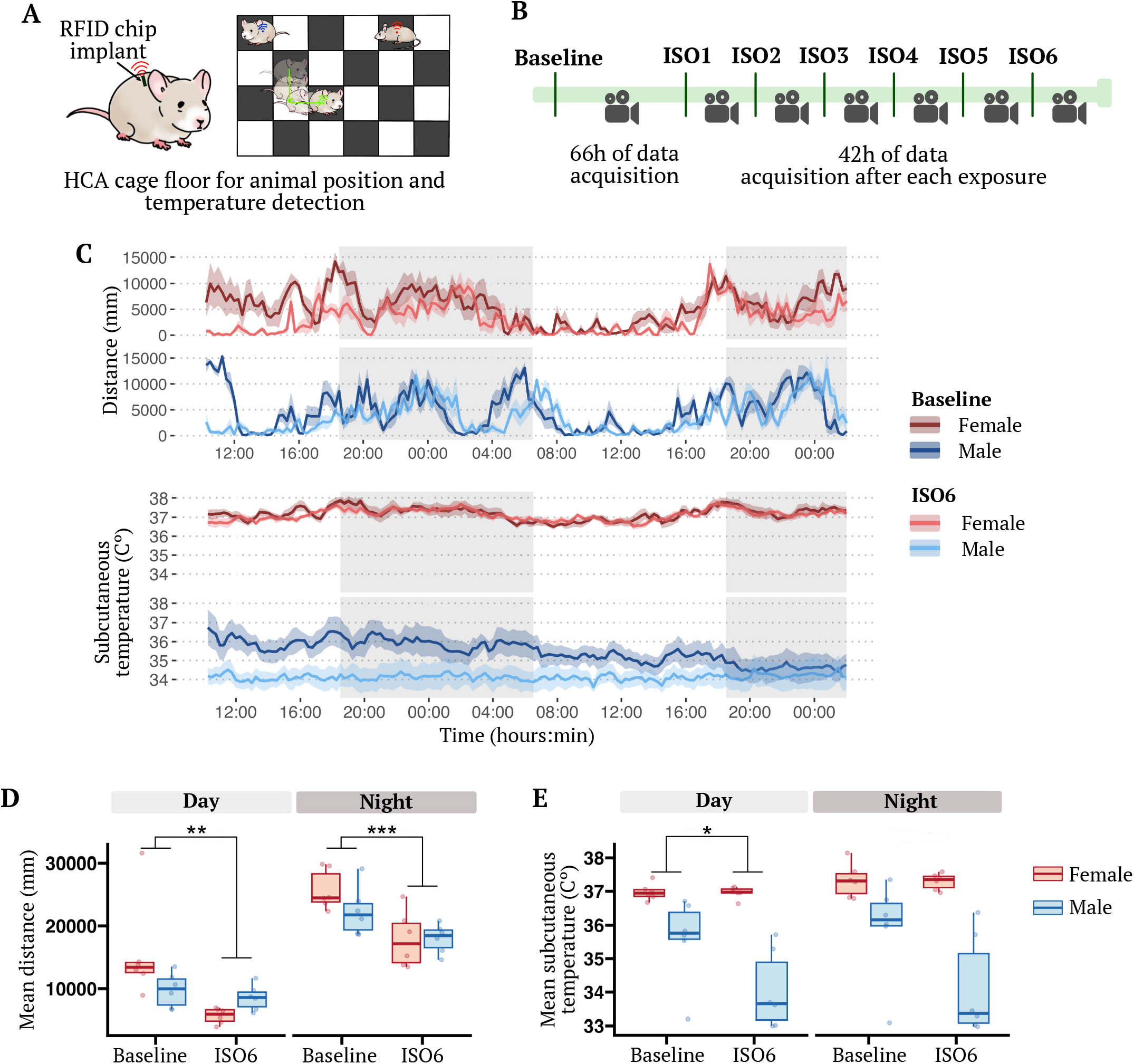
Effects of repeated isoflurane induction on spontaneous motor activity and subcutaneous temperature in mice as measured by automated home-cage analysis. A) Schematic depiction of the home-cage analysis (HCA) system used for continuous monitoring of animal position, locomotor activity, and subcutaneous temperature via implanted RFID chips. B) Experimental design schematic showing timeline of baseline recording, repeated isoflurane induction sessions (ISO1–ISO6) and post-anaesthesia recording periods. C) Typical cage-level recordings of baseline and post-repeated isoflurane (ISO6) recordings of mean distance travelled (top) and mean subcutaneous temperature (bottom) over the 42 h recording period across consecutive day and night (grey shaded) cycles for female (red/coral) and male (navy/blue) animals; data are mean ± SEM, n=6 (3 mice/cage). D) Mean distance travelled during day and night periods for female (red) and male (blue) mice at baseline and for the 42 h period following the sixth cycle of brief isoflurane anaesthesia; data are mean ± SEM, n=6 mice/ group, ** P<0.01, *** P<0.001. E) Mean subcutaneous temperature during day and night periods for female (red) and male (blue) mice at baseline and for the 42 h period following the sixth cycle of brief isoflurane anaesthesia; data are mean ± SEM, n=6 mice/ group, ** P<0.01, *** P<0.001.

Body temperature was influenced by repeated isoflurane treatment during the light phase (F_1,20_=4.742, P=0.042), and pronounced sex differences were observed in both the light (F_1,20_=36.135, P<0.001) and dark (F_1,20_=26.305, P<0.001) phases. Male animals exhibited a consistent reduction in mean body temperature following anaesthetic treatment (light phase P=0.03; dark phase P=0.048; Figure 1E), while female mice maintained comparable temperatures at baseline and after exposure (Figure 1E). The post-mortem assessment of the RFID chips positions confirmed the subcutaneous location in the abdomen, with this location being generally lower in the male animals. These temperature readouts thus may be more associated with peripheral rather than core body temperature.

### Repeated anaesthesia alters expression of murine maintenance behaviours

Mice exhibit numerous behaviours beyond locomotor activity; hence we employed a behavioural ethogram (Supplemental table 1) to assess the impact of repeated anaesthetic exposure upon key maintenance, exploratory and social behaviours, again within their home cage environment, recorded over two thirty-minute windows across the transitions from light to dark and dark to light phases, i.e. 15 minutes prior to and after lights turning on or off respectively (Figure 2A).

**Figure 2:**
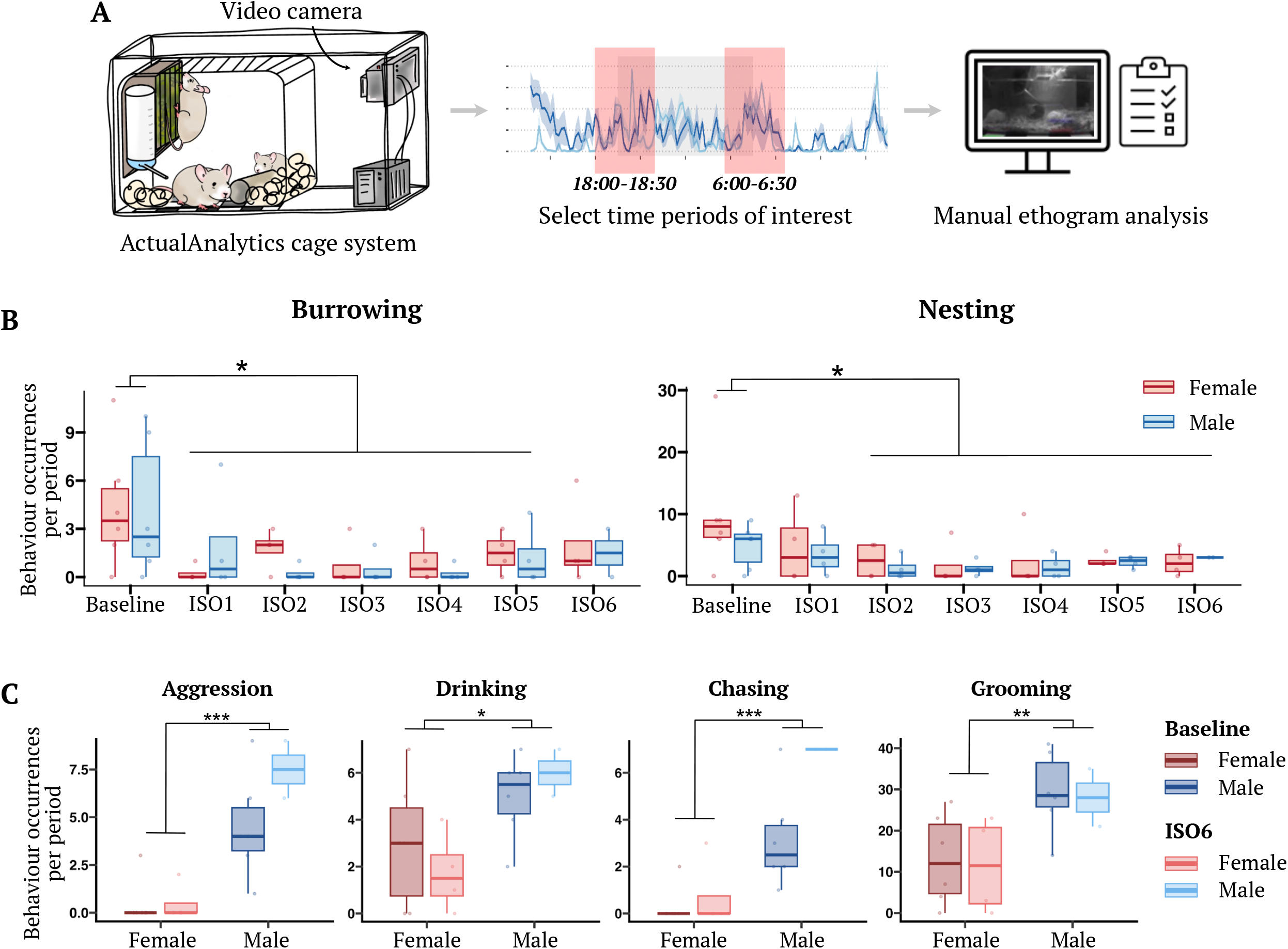
Spontaneous burrowing and nesting behaviours are suppressed following repeated isoflurane induction. A) Schematic representation of the protocol for manual analysis of spontaneous behaviours during baseline conditions and following exposure to repeated isoflurane anaesthesia. B) Frequency of burrowing (left) and nesting (right) behaviours per observation period during baseline and following each isoflurane induction session (ISO1–ISO6) for female (red) and male (blue) mice; data are mean ± SEM, n=2-6 group-housed cages of 3 mice, *P<0.05. C) Frequency of aggression, drinking, chasing, and grooming behaviours per observation period during baseline and post-isoflurane treatment for female (red) and male (blue) mice; no effects of isoflurane were apparent at any point; data are mean ± SEM, n=2-6 group-housed cages of 3 mice, *P<0.05, **P<0.01, ***P<0.01.

As might be expected, clear effects of time of day were seen in many behaviours, with substantially increased incidences of feeding (F_1,26_=14.84, P<0.001), rearing (<3 s: F_1,26_=7.96, P=0.009; ≥3 s: F_1,26_=17.03, P<0.001), climbing (F_1,26_=16.012, P<0.001), hanging (F_1,26_=15.743, P<0.001), and social interactions such as anogenital investigation (F_1,26_=9.42, P=0.005) and allo-grooming (F_1,26_=4.871, P=0.036) during the period of transition to wakefulness (Supplemental Table 2). In some cases, sex differences were apparent in behaviour frequency, with male dominant effects seen in in chasing (F_1,26_=9.193, P=0.005) and aggressive behaviours (F_1,26_=12.617, P=0.001), and sex by time of day interactions seen in grooming (F_1,26_=7.198, P=0.013) and scrape digging (F_1,26_=4.233, P=0.049), where in both cases females exhibited lower frequencies of these behaviours in the morning than the evening, although males showed no temporal differences; the opposite pattern was seen in tube running (F_1,26_=5.819, P=0.023), i.e. males were more active in the mornings while females showed similar levels of behaviour regardless of time of day.

Two main effects of repeated anaesthetic exposure upon behavioural frequency were apparent, affecting nesting and burrowing behaviours; in neither case were sex differences or significant interactions with the time of day when measured apparent. For nesting behaviours, animals exposed to isoflurane showed significantly and persistently lower frequencies after the second round of isoflurane exposure (F_6,26_=3.586, P=0.01), and animals showed reduced incidences of burrowing behaviour from the first exposure to isoflurane through to the fifth round (F_6,26_=2.706, P=0.035), although statistical significance on this measure was lost by the sixth and final isoflurane exposure.

### Sex differences in cerebral blood flow reactivity are enhanced following repeated anaesthetic exposure

Anaesthesia is known to affect cerebral blood flow both directly and secondarily following changes in brain cell function (Slupe and Kirsch, 2018), hence we examined whether repeated isoflurane exposure would affect surface brain blood flow using laser speckle perfusion imaging, assessed both under standard anaesthesia with 2% isoflurane in oxygen and following two separate, transient challenges of increased isoflurane dose, increasing to 5% isoflurane in oxygen for one minute (Figure 3A-B) on two occasions five minutes apart. No significant differences were apparent between control animals and those that had been exposed to repeated isoflurane treatment in either baseline cerebral perfusion or following 5% isoflurane challenge, although females tended to exhibit greater cerebral perfusion than males, although this did not reach statistical significance (F_1,13_=4.593 P=0.052). Repeated challenge with 5% isoflurane resulted in a decline in cerebral perfusion in all cases (F_1,13_=31.035, P<0.001), with males exhibiting a greater decrease in cerebral perfusion than females irrespective of trial (F_1,13_=56.674, p=0.023), suggestive of functional sex differences within the cerebral vasculature; again this was unaffected by previous repeated isoflurane anaesthesia (Figure 3C). Reaction time to the isoflurane challenge was also examined to determine whether repeated anaesthetic exposure altered responsiveness to the 5% isoflurane challenge. Interestingly, sex had an effect on reaction time, with male animals responding faster overall than females (P=0.039), independent of treatment condition or trial. In contrast, there was no significant main effect of isoflurane treatment, nor evidence for treatment-by-sex or higher-order interactions, indicating that repeated anaesthetic exposure did not alter reaction time to the anaesthetic challenge (Figure 3D).

**Figure 3:**
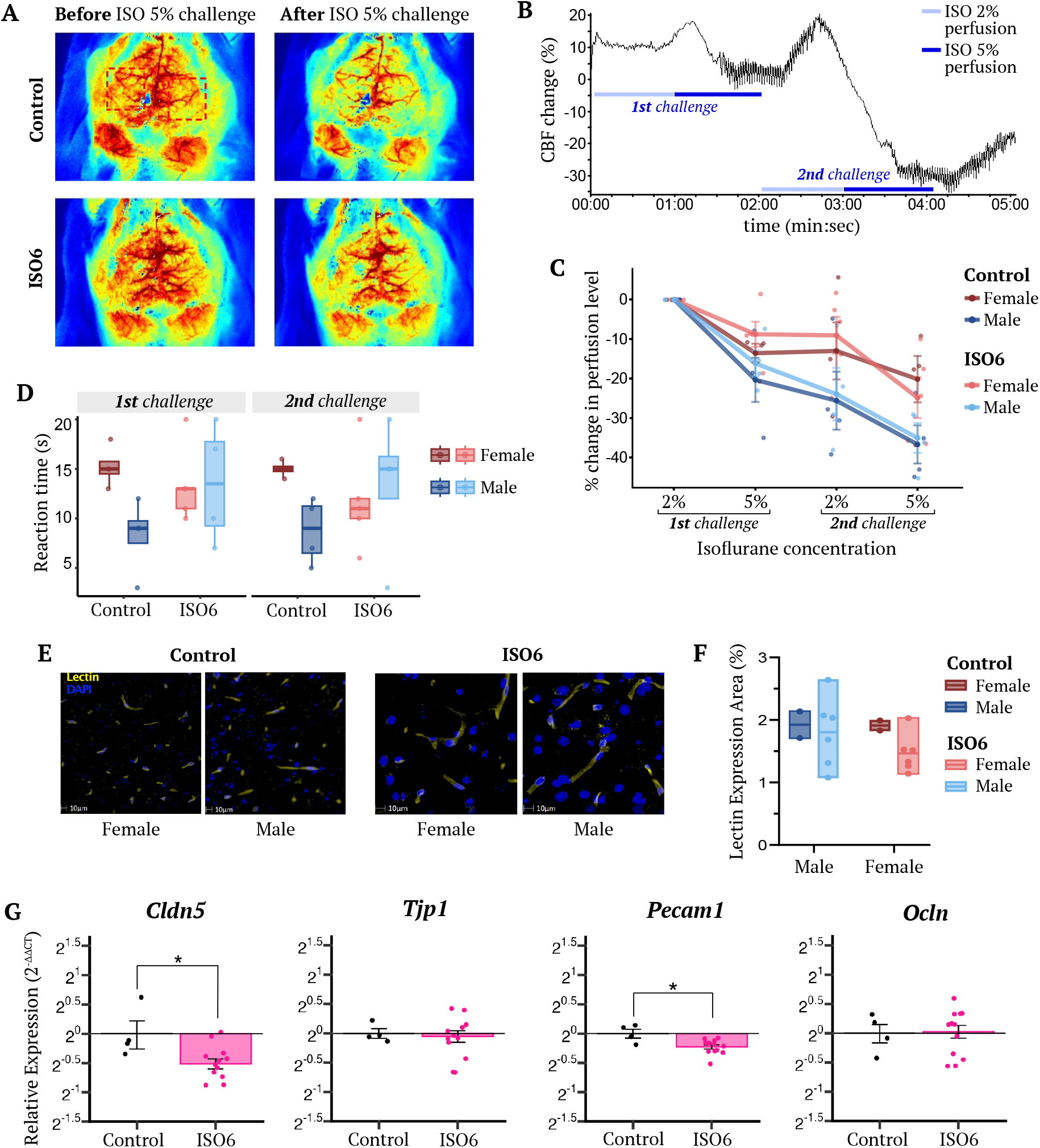
Repeated isoflurane exposure alters molecular components of the cerebral vasculature without significant effects on cerebral blood perfusion rate or reactivity. A) Representative cerebral blood perfusion images showing perfusion at baseline and following repeated isoflurane anaesthesia before and after exposure to a 5% isoflurane challenge. For all analyses, perfusion values from regions of interest (ROIs) in the left and right hemispheres were averaged. B) Representative real time cerebral blood flow trace indicating the time windows corresponding to 2% (pink) and 5% (red) isoflurane exposure. C) Cerebral blood perfusion during repeated periods of 2% and 5% isoflurane exposure, expressed as percentage change from mean baseline perfusion prior to the first challenge, for female (red/coral) and male (navy/blue) mice at baseline and post-repeated isoflurane exposure; data are mean ± SEM, n=4 mice/group. D) Time taken for cerebral blood flow to stabilise following administration of a 5% isoflurane challenge, for female (red/pink) and male (blue/cyan) mice at baseline and post-repeated isoflurane exposure; data are mean ± SEM, n=4 mice/group. E) Representative images of cerebral vascular labelling using AF488-conjugated tomato lectin (yellow), with nuclear counterstaining using DAPI (blue) in the cerebral cortex of naïve mice and post–repeated isoflurane exposure; sections are 10 μm thickness, scale bar is 10 μm. F) Lectin-positive vascular area (%) in naïve mice and following repeated isoflurane exposure mice (left) and stratified by sex (right); data are mean ± SEM, n=6 mice/group (n=4 control) . G) Relative whole brain mRNA expression for vascular-associated genes (*Cldn5, Tjp1, Ocln, Pecam1*) in naïve mice and following repeated isoflurane exposure; data are mean ± SEM, n=6-12 mice, *P<0.05.

To understand whether these changes in cerebral perfusion reflected underlying changes in vascular density, we examined blood vessel density within the cortical parenchyma. We saw no significant effects of repeated isoflurane treatment (Figure 3D-E). However, RT-qPCR analysis of the expression of key vascular genes revealed a significant down-regulation in mRNA for claudin-5 (P=0.019) and platelet endothelial cell adhesion molecule-1 (PECAM-1) (P=0.016), though not the other endothelial tight junction components occludin or zonula occludens-1 (Figure 3F).

### Repeated isoflurane treatment suppresses pro-inflammatory microglial markers

Given the well-established links between post-operative delirium and neuroinflammation (Li et al., 2025), we examined whether repeated isoflurane exposure affected key neuroinflammatory parameters, namely cytokine expression and microglial phenotype. Contrary to our expectations, we saw no evidence for ongoing neuroinflammatory change, with no changes in mRNA expression for either the pro-inflammatory cytokines tumour necrosis factor (TNF)α, interleukin (IL)-1β or for the anti-inflammatory cytokine IL-10 (Figure 4A), nor any sign of neuronal/synaptic damage, as assessed by measuring expression of the synaptic components synaptophysin and PSD-95 (Figure 4B). However, examination of microglial phenotype revealed a significant reduction in expression of the major pro-inflammatory markers CD40 (F_1,28_=10.04, P=0.004) and CD68 (F_1,28_=14.971, P<0.001) in male and female animals exposed to isoflurane, when compared with sham treated controls (Figure 4C), indicative of a hyporesponsive microglial status.

**Figure 4:**
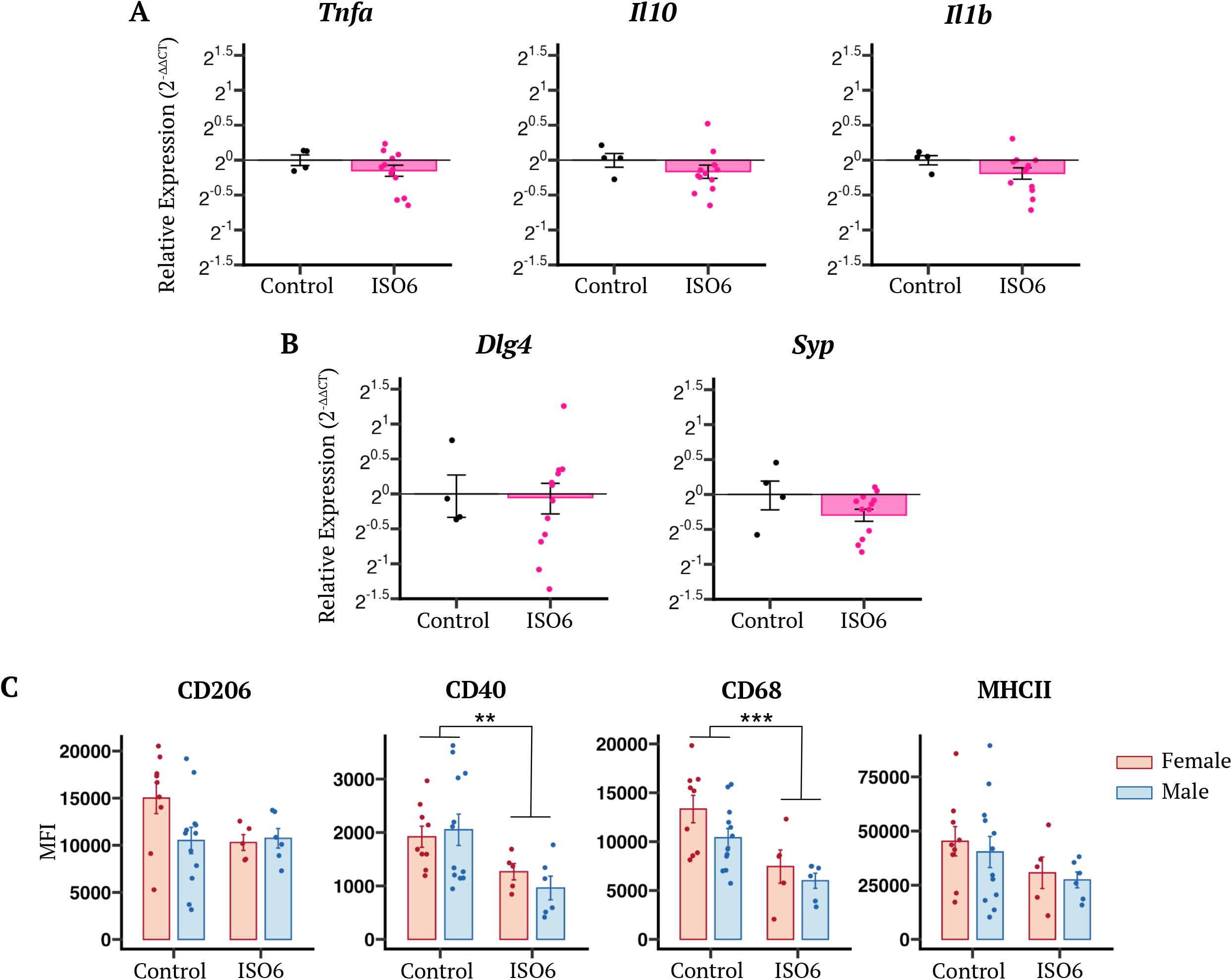
Repeated isoflurane exposure suppresses key microglial activation markers without affecting gene expression of cytokines or synaptic markers. A) Relative whole brain mRNA expression of pro- and anti-inflammatory cytokine genes (*Tnfa, Il1b, Il10*) in naïve mice and following repeated isoflurane exposure; data are mean ± SEM, n=4-12 mice. B) Relative whole brain mRNA expression of synaptic genes (*Dlg4, Syp1*) in naïve mice and following repeated isoflurane exposure; data are mean ± SEM, n=24 mice. C) Cell surface expression of phenotypic markers (CD206, CD40, CD68, MHCII) on isolated microglial cells from naïve mice and those exposed to repeated isoflurane anaesthesia, expressed as median fluorescence intensity (MFI); data are mean ± SEM, n=6-12 mice, **P<0.01, ***P<0.001.

## Discussion

Exposure to general anaesthetics has been linked to cognitive disruption in both clinical research and in pre-clinical models, but whether repeated rounds of anaesthesia and recovery, as are commonly used in animal studies, have similar effects is less well established (Walton, 1979; Schneemilch et al., 2004; Hohlbaum et al., 2017; Bajwa et al., 2019). In this study, we demonstrate that repeated short isoflurane exposures induce subtle but measurable changes in behaviour and microglial phenotype, with several of these outcomes showing clear sex specificity. Together, we propose, these data highlight the importance of including anaesthetic regime as a relevant variable when analysing the outcome of experimental models, particularly those related to the brain and its functions.

Behaviourally, repeated isoflurane exposure suppressed total locomotor activity in a sex-selective manner. Female mice exhibited reduced locomotor activity across both light and dark phases, whereas males did not, aligning with previous studies reporting females to be more susceptible to isoflurane-induced motor alterations (Hohlbaum et al., 2017; Bajwa et al., 2019). In contrast, both sexes displayed diminished nesting and burrowing behaviour, two ethologically important maintenance behaviours that reflect welfare, motivation, and stress coping (Jirkof, 2014; Gjendal et al., 2019; Domínguez-Oliva et al., 2025). Prior research has documented mixed effects of isoflurane on these behaviours, with some studies reporting burrowing impairments only and others reporting effects of acute but not repeated isoflurane treatment on nesting (Hohlbaum et al., 2017; Gjendal et al., 2019). This difference may largely be attributable to the differing dosing regimens used, i.e., 15 minutes of isoflurane repeated several weeks apart in the earlier work versus 5 minutes every 2-3 days in our study. Nonetheless, these studies together serve to highlight both that isoflurane anaesthesia can influence major behavioural parameters, across different species, and that this is not always fully predictable, emphasising the need for caution and consideration in the use of anaesthesia in the development of preclinical models, particularly those that will require behavioural analysis.

While our study found no evidence to indicate isoflurane-induced changes in the markers of synaptic density, PSD-95 and synaptophysin, there remain several plausible pathways through which isoflurane may influence neuronal function. Isoflurane acutely modulates neurotransmitters, including acetylcholine, dopamine, and noradrenaline in the prefrontal cortex (Zhang et al., 2020), a region implicated in goal-directed and maintenance behaviours such as nesting and burrowing (Deacon et al., 2003). We have also shown that repeated exposure to isoflurane activates a cellular senescence response in immature mice which could also be linked to subtle behavioral disturbances (Al-Khateeb et al., 2024). Isoflurane also activates the stress response, with females showing stronger corticosterone responses in some studies (Deckardt et al., 2007; Wu et al., 2015; Bekhbat et al., 2016), which may contribute to the observed sex-biased locomotor suppression. Fuller investigation of these, or other, potential mechanistic targets will be required in future, but they do begin to suggest how repeated anaesthetic exposures could lead to persistent behavioural effects.

While cerebrovascular function did not seem to be affected by repeated isoflurane exposure, there were signs of sex differences in several aspects. Across both baseline conditions and following repeated isoflurane exposure, females exhibited an elevated superficial cerebral blood flow compared to males; however, these differences did not reach statistical significance. Interestingly, while both females and males exhibited a decline in cerebral perfusion in response to 5% isoflurane challenge, this was significantly greater in males, suggesting intrinsic differences in cerebrovascular reactivity. Prior work has identified sexual dimorphisms in cerebrovascular architecture (Maggioli et al., 2016; Collignon et al., 2024) and has shown that isoflurane interacts with estrogen signalling in complex ways, with isoflurane preconditioning being shown to protect against cerebral ischaemia in male mice, be neutral in older females, and be detrimental in young female mice (Kitano et al., 2007; Wang et al., 2008). Our findings build on this literature by demonstrating subtle sex differences in cerebrovascular function and reactivity, particularly in response to challenge. It is clear that more work is needed to unravel these interactions, but they serve to highlight the absolute need to consider sex as an experimental variable in preclinical and clinical studies.

While repeated isoflurane exposure did not cause any obvious alterations in cortical capillary density, reduced expression of endothelial markers claudin-5 and PECAM-1 were evident, suggesting molecular rather than anatomical modifications to the vasculature. PECAM-1 has been shown to be involved in the response to altered cerebral blood flow (Chen and Tzima, 2009), raising the question of whether more subtle changes in perfusion dynamics at the level of individual blood vessels may have occurred following repeated isoflurane exposure. Claudin-5 is preferentially expressed in cerebral capillaries over larger blood vessels in the brain (Hashimoto et al., 2023), again suggesting that localised changes to cerebrovascular function may have occurred in response to repeated isoflurane treatment. Notably, it is well established that extended periods of isoflurane treatment can cause vasodilation (Oshima et al., 2003; Aksenov et al., 2015; Sullender et al., 2022), driven in part by direct influences of the anaesthetic on endothelial biology, including reduced claudin-5 expression and consequent destabilisation of tight-junction organisation (Thal et al., 2012; Zhang et al., 2021), and by impaired nitric oxide (NO) signalling and mitochondrial stress (Noorani et al., 2023). In contrast, the impact of repeated brief isoflurane exposures on long-term cerebrovascular parameters has received limited attention; our data provide initial insights into such changes.

Finally, repeated exposure influenced microglial phenotype, with both sexes showing decreased expression of activation-associated markers CD40 and CD68, although both pro- and anti-inflammatory cytokine levels remained unchanged. This pattern suggests a shift toward a less immunoresponsive microglial state (Tan et al., 1999; Swanson et al., 2023), contrasting with previous studies using prolonged (hours-long) isoflurane exposures, which often report pro-inflammatory effects, including activation of NFκB (Cao et al., 2018), disruption of astrocyte gap junctions (Dong et al., 2022) and induction of microgliosis and production of the pro-inflammatory cytokines TNFα, IL-1β and IL-6 (Bai and Shi, 2025). Taken together, this indicates that immune outcomes are highly dependent on exposure duration and frequency, with brief repeated exposures producing a distinct profile.

This study is not without limitations, most prominently in that we used a relatively brief, albeit repeated, exposure to isoflurane of five minutes, and only one strain of mice, CD-1. Longer durations of treatment, and extension to different mouse lines, may produce quantitatively or qualitatively different effects, as was seen in studies of the effects of repeated isoflurane on nesting behaviour for example (Gjendal et al., 2019). Nonetheless, the pattern of anaesthesia we have employed reflects those commonly used in, e.g., longitudinal imaging studies, but further work will be required to extend these findings to different strains of mouse, and to consider the interactions of anaesthesia with factors such as age and genotype, broadening its relevance. Further mechanistic studies are needed to elucidate these adaptive effects observed after repeated exposure, particularly given isoflurane’s complex and multifaceted pharmacodynamics and multiple specific cellular targets.

Repeated short anaesthetic exposures, although commonplace and often considered minimal to non-invasive methods, constitute a biologically active intervention capable of shaping brain homeostasis. Our findings demonstrate that repeated isoflurane exposure produces subtle but coordinated shifts in behaviour, cerebrovascular function, and microglial activation, several of which differ between males and females. Although anaesthesia is essential for ensuring animal welfare, its effects are not physiologically neutral, and its cumulative effects have the potential to confound interpretations of neurological and behavioural experiments. These observations highlight the need to treat the anaesthetic regimen as a meaningful experimental variable, deserving the same attention and scrutiny as other procedural factors. For researchers, this means minimising non-essential exposures, ensuring appropriate recovery periods, incorporating precisely matched sham-anaesthetised controls, and considering sex differences in both study design and analysis. From an animal welfare perspective, the fact that even brief routine isoflurane inductions can influence behaviour and physiology underscores the importance of refining protocols, ensuring physiological monitoring and supportive approaches to maintain tissue homeostasis, preserving hemodynamic stability and oxygenation, and temperature regulation to reduce cumulative burden, while supporting good recovery from any deviations in physiological variables. Together, these considerations will be critical for improving the reproducibility, and translational relevance of preclinical studies that involve repeated anaesthesia.

## Acknowledgements

We would like to thank Brian Lock and Moor Instruments Ltd, for the use of the moorFLPI Full-Field Laser Perfusion Imager and for their support in the cerebral perfusion studies, Rowland Sillito and Actual Analytics Ltd, UK for their guidance and technical support with the HCA system, the excellent team of animal technologists at QMUL Biological Services Unit for their animal husbandry and care. We thank the Neuroscience Educational teams at QMUL for their financial support to enable students to engage with these research programmes; in particular, Dr Ping Yip for his unconditional collegiality and support with these projects.

## Author contributions

JLT, ZS and TR, carried out the behaviour studies; SMcA, IV and SHK carried out the flow cytometry and qPCR studies; JLT and SMcA carried out the CBF studies; JLT and OM carried out the IHC studies, ZS and SMcA carried out data analysis; ZS, JLT, SMcA wrote the manuscript; JLT and SMcA designed the study and managed the funding; ZS, JLT, SMcA, AMT reviewed the manuscript.

## Figure Legends

**Supplemental Figure 1:**
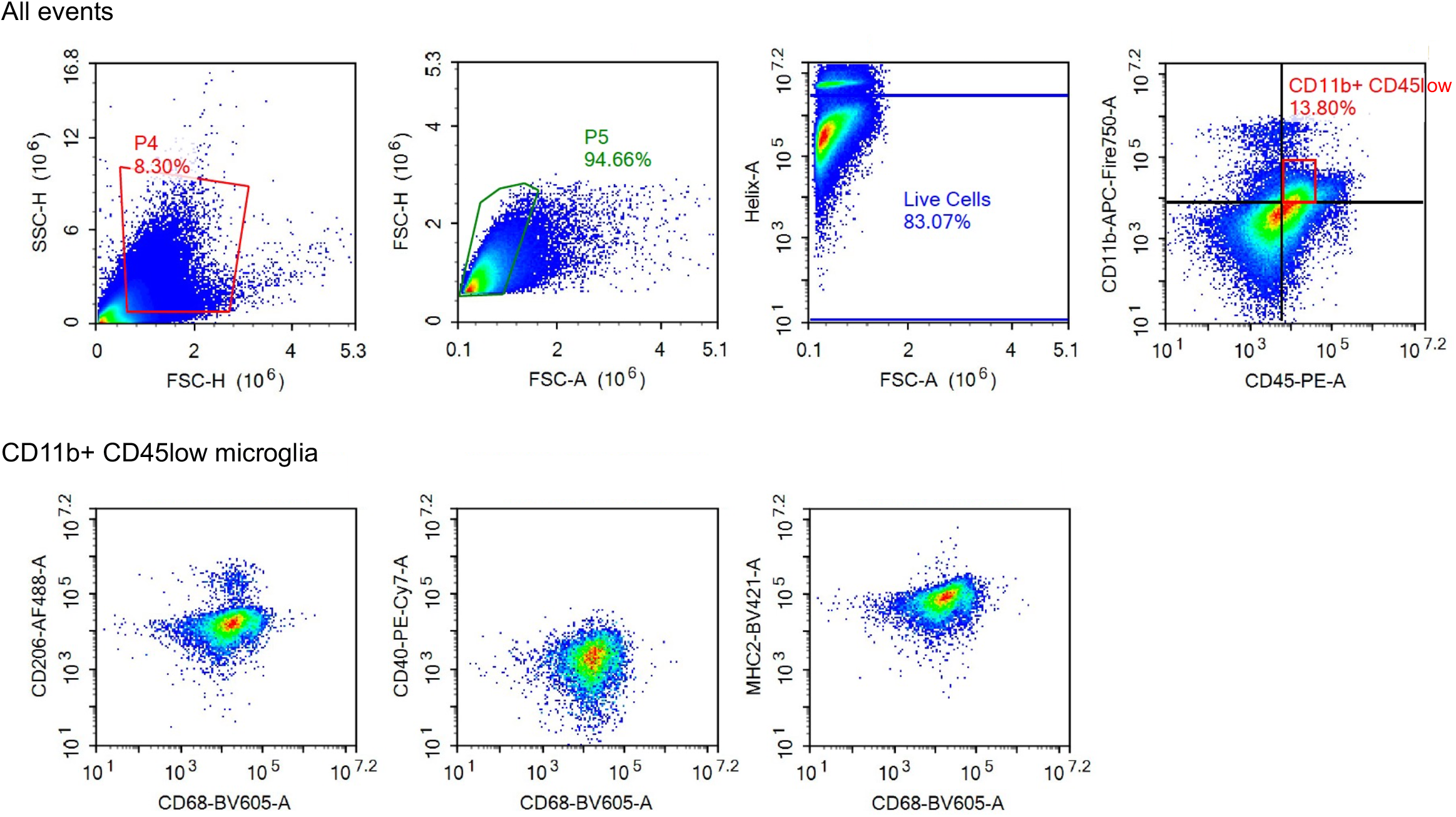
Gating strategy for identification of primary microglia (CD45 low, CD11b low) and typical scatterplots for phenotypic markers (CD68, CD206, CD40 & MHCII).

**Supplemental Table 1:**
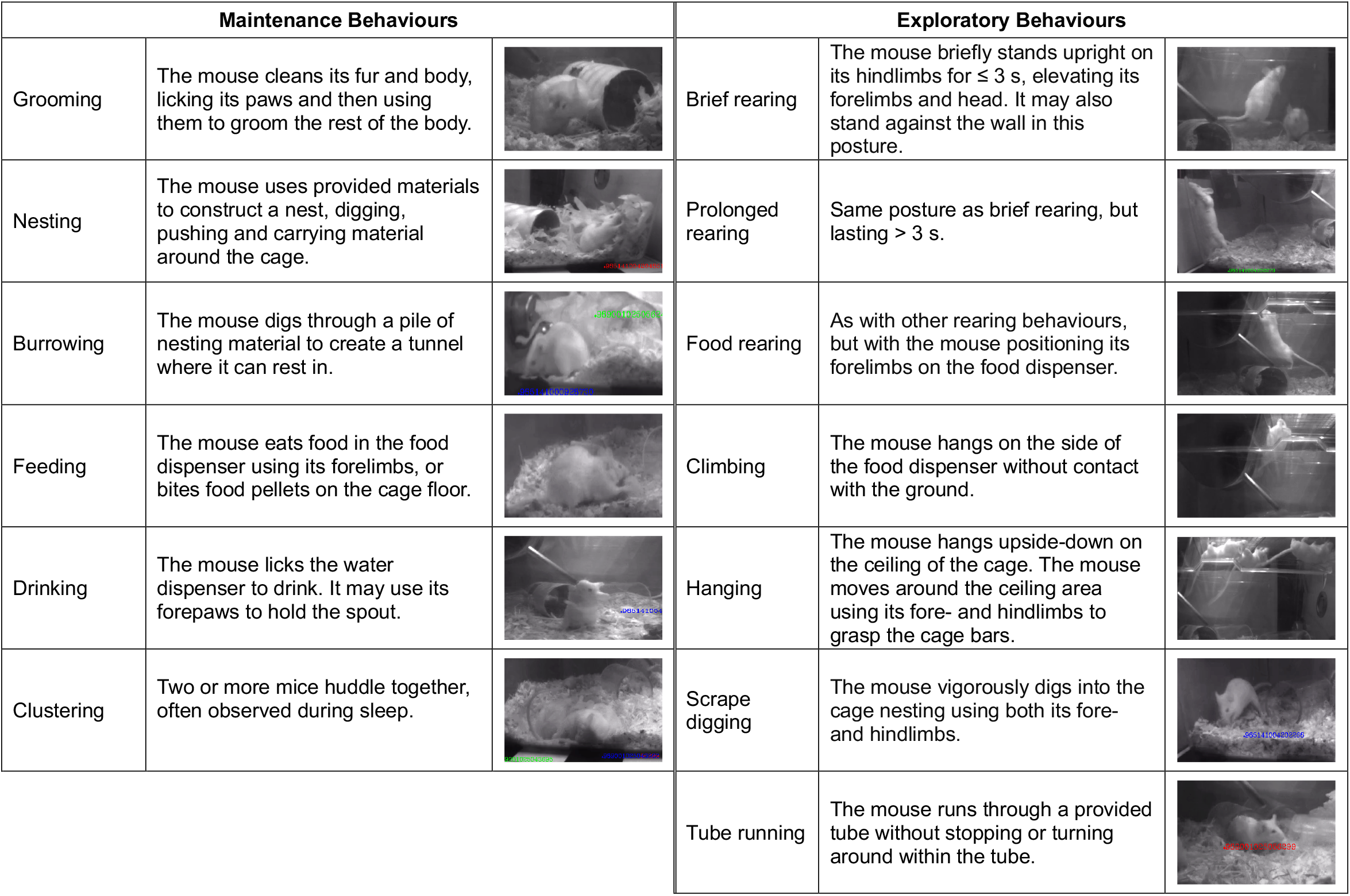
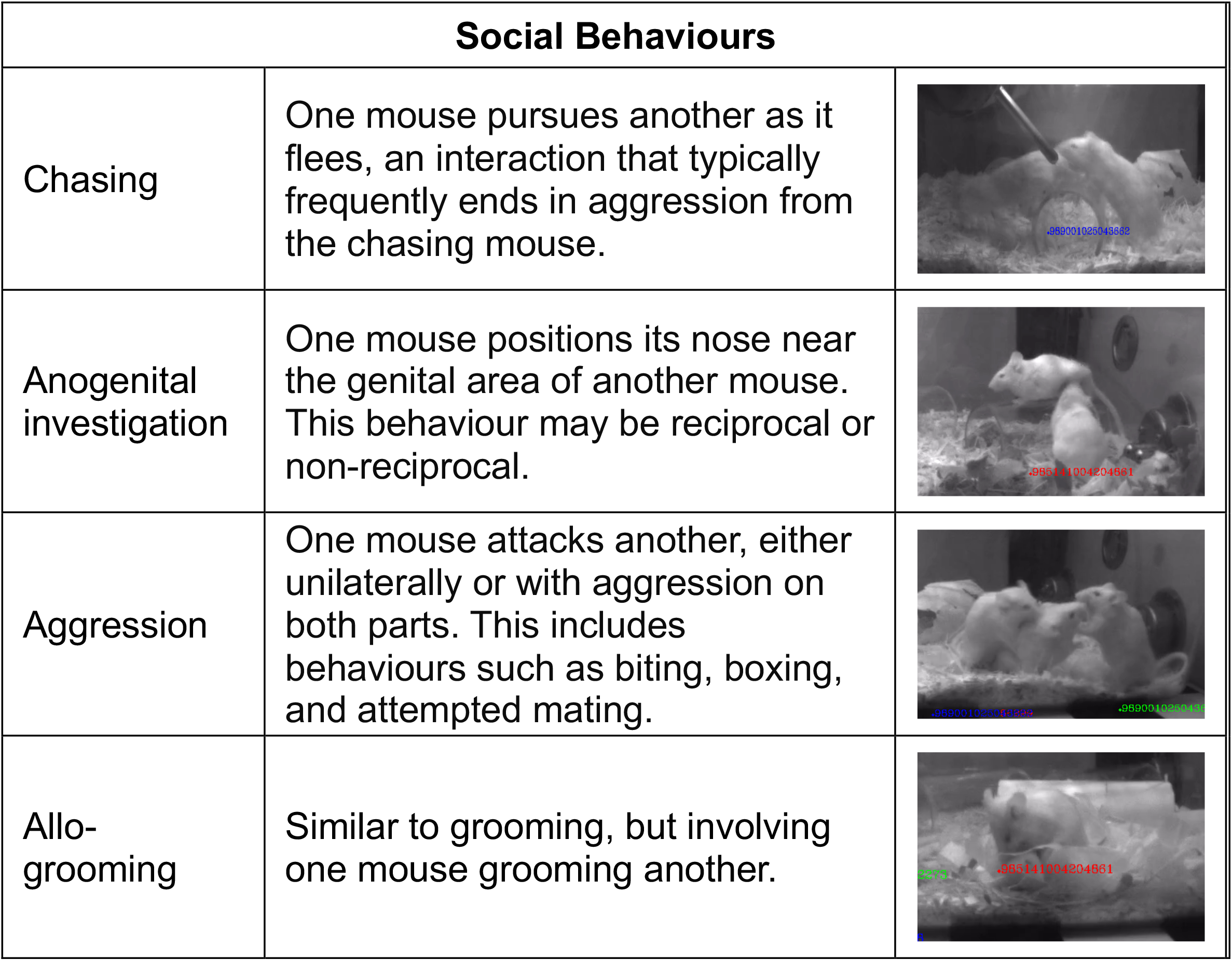
Ethogram depicting typical murine behaviours analysed in infrared video recordings, grouped as maintenance, exploratory or social behaviours.

| Maintenance Behaviours |  |  | Exploratory Behaviours |  |  |
| --- | --- | --- | --- | --- | --- |
| Grooming | The mouse cleans its fur and body, licking its paws and then using them to groom the rest of the body. | | Brief rearing | The mouse briefly stands upright on its hindlimbs for $\leq 3$ s, elevating its forelimbs and head. It may also stand against the wall in this posture. | |
| Nesting | The mouse uses provided materials to construct a nest, digging, pushing and carrying material around the cage. | | Prolonged rearing | Same posture as brief rearing, but lasting $> 3$ s. | |
| Burrowing | The mouse digs through a pile of nesting material to create a tunnel where it can rest in. |  | Food rearing | As with other rearing behaviours, but with the mouse positioning its forelimbs on the food dispenser. |  |
| Feeding | The mouse eats food in the food dispenser using its forelimbs, or bites food pellets on the cage floor. |  | Climbing | The mouse hangs on the side of the food dispenser without contact with the ground. |  |
| Drinking | The mouse licks the water dispenser to drink. It may use its forepaws to hold the spout. |  | Hanging | The mouse hangs upside-down on the ceiling of the cage. The mouse moves around the ceiling area using its fore- and hindlimbs to grasp the cage bars. |  |
| Clustering | Two or more mice huddle together, often observed during sleep. |  | Scrape digging | The mouse vigorously digs into the cage nesting using both its fore- and hindlimbs. |  |
|  |  |  | Tube running | The mouse runs through a provided tube without stopping or turning around within the tube. |  |

| Social Behaviours |  |
| --- | --- |
| Chasing | One mouse pursues another as it flees, an interaction that typically frequently ends in aggression from the chasing mouse. |
| Anogenital investigation | One mouse positions its nose near the genital area of another mouse. This behaviour may be reciprocal or non-reciprocal. |
| Aggression | One mouse attacks another, either unilaterally or with aggression on both parts. This includes behaviours such as biting, boxing, and attempted mating. |
| Allo-grooming | Similar to grooming, but involving one mouse grooming another. |

